# Spectral and lifetime fluorescence unmixing via deep learning

**DOI:** 10.1101/745216

**Authors:** Jason T. Smith, Marien Ochoa, Xavier R. M. Intes

## Abstract

Hyperspectral Fluorescence Lifetime Imaging allows for the simultaneous acquisition of spectrally resolved temporal fluorescence emission decays. In turn, the acquired rich multidimensional data set enables simultaneous imaging of multiple fluorescent species for a comprehensive molecular assessment of biotissues. However, to enable quantitative imaging, inherent spectral overlap between the considered fluorescent probes and potential bleed-through must be taken into account. Such task is performed via either spectral or lifetime unmixing, typically independently. Herein, we present UNMIX-ME (unmix multiple emissions), a deep learning-based fluorescence unmixing routine, capable of quantitative fluorophore unmixing by simultaneously using both spectral and temporal signatures. UNMIX-ME was trained and validated using an *in silico* framework replicating the data acquisition process of a compressive hyperspectral fluorescent lifetime imaging platform (HMFLI). It was benchmarked against a conventional LSQ method for both tri and quadri-exponential simulated samples. Last, UNMIX-ME’s potential was assessed for NIR FRET *in vitro* and *in vivo* for small animal experimental data.

---

Fluorescence imaging is the most employed molecular imaging technique from the wet lab to the bed side. A key strength of fluorescence imaging is its ability to simultaneously image multiple fluorophores (multiplexing) for improved understanding of the sample’s molecular features. Typically, multiplexing is achieved via selection of exogenous fluorophores with distinct spectral features. Though, spectral overlap of the fluorophore’s emission is unavoidable, leading to bleed-through between acquisition channels. This is also an outstanding challenge in endogenous imaging, in which multiple species can be simultaneously excited at a given wavelength. Hence, spectral imaging is always associated with spectral unmixing algorithms that leverage the spectral signature of each individual fluorophore to determine their individual contributions at each pixel from the raw fluorescence images. Such unmixing methodologies are often based on *a priori* fitting techniques that use publicly available or experimentally acquired “pure” spectra [1]. Though, spectral imaging and linear unmixing are sensitive to noise, large spectral overlap and/or wrong or incomplete spectral information [2]. Besides fluorescence intensity, it is also possible with dedicated instruments to quantify fluorescence lifetime, which is an intrinsic characteristic of fluorophores. However, for biomedical applications, it is challenging to perform unmixing beyond two lifetime contributions due to the low level of lights typically encountered. Recently, there has been great interest in performing multi- or hyper-spectral Fluorescence Lifetime Imaging (FLI) to augment the potential of lifetime imaging for highly multiplexed studies [3]. Especially, coupling spectral unmixing with FLI has the potential to achieve significantly higher unmixing sensitivity and specificity than that of intensity-based or lifetime-based methods alone. [4] Despite great progress in instrumentation that helps collect such multidimensional data sets, the approach to perform unmixing is still typically applied along spectra or time, not both dimensions together. Herein, we propose “UNMIX-ME” (unmix multiple emissions), a deep convolutional neural network (CNN) -based framework that performs fluorophore unmixing by leveraging both spectral and lifetime contrast concomitantly.

UNMIX-ME is designed to retrieve the spatially resolved abundance coefficients associated with the sample’s fluorophores from the spectrally resolved fluorescence decay measurements. The proposed methodology is developed within the context of Hyperspectral Macroscopic Fluorescence Lifetime Imaging (HMFLI). Such approach is based on a recently proposed novel instrumental concept that leverages a single-pixel strategy to concurrently acquire 16 spectrally resolved FLI channels over large field of views (FOV). [3] Through the use of Deep Learning (DL), HMFLI has proven capable of probing nanoscale biomolecular interactions across large fields of view (FOV) at resolutions as high as 128×128 within minutes. [5] DL has also greatly improved the processing time for its inverse solving procedure, yielding intensity and lifetime reconstructions in a single framework, through usage of simulated training data mimicking the single-pixel data generation. [6] UNMIX-ME aims to further enhance the HMFLI hyperspectral toolbox by facilitating accurate unmixing capabilities. First, we report on the design of UNMIX-ME architecture. Then we describe the novel data simulation routine developed to efficiently generate 16-channel fluorescence temporal point-spread functions (TPSFs) used to train our CNN – which bypasses the need of collecting large quantities of experimental data and enables the enforcement of correct parametric mapping to ground-truth instead of relying on fitting procedures. The performances of UNMIX-ME are reported for both the case of tri and quadri-component unmixing *in silico*. To further validate UNMIX-ME, we report on its capability to process *in vitro* HMFLI data sets of Föster Resonance Energy Transfer (FRET) with excitation and emission spectra in the near-infrared (NIR) – i.e. fluorophores currently known to possess short lifetime values (sub-nanosecond) and correspondingly high analytic complexity. Lastly, we present hyperspectral lifetime unmixing results for two *in vivo* datasets as acquired in [9]: 1) Trastuzumab (TZM) AF700/AF750-conjugated FRET pair, for an athymic nude mouse bearing a tumor xenograft and imaged 76 hours post-injection and 2) Transferrin (Tf) AF700/AF750-conjugated FRET pair to distinguish between mouse liver and bladder.

UNMIX-ME’s CNN architecture (Fig. 1) was crafted such that extraction of temporal information was prioritized while mitigating the computational burden associated with processing simultaneously 16 spectral channels-worth of TPSF data. Given that the use of 3D convolutional operations (Conv3D) is notoriously expensive computationally, just two Conv3D layers employing large stride were included – allowing for significant reduction in parametric size within the early layers. The output from these layers was transformed into 2D and followed by 2D separable convolutions [8] with kernel size 1×1 as a more computationally friendly alternative for spatially-independent temporal and spectral feature extraction. Moreover, “XceptionBlock” [7] operations (i.e., residual blocks with 1×1 separable convolutions) were included to ensure that our model would reap the benefits obtained through residual learning [10] while maintaining focus on the primary objective – spatially-independent temporal and spectral feature extraction. UNMIX-ME introduces the concept of “CoefficientBlocks”, individual branches composed of a set of 2D convolutions intercepted by batch normalization and activation layers, with each branch meant to focus on features relevant for fluorophore-specific abundance coefficient retrieval. These blocks facilitate seamless architecture modification for retrieval of *N* number of targeted fluorophores.

**Fig 1.**
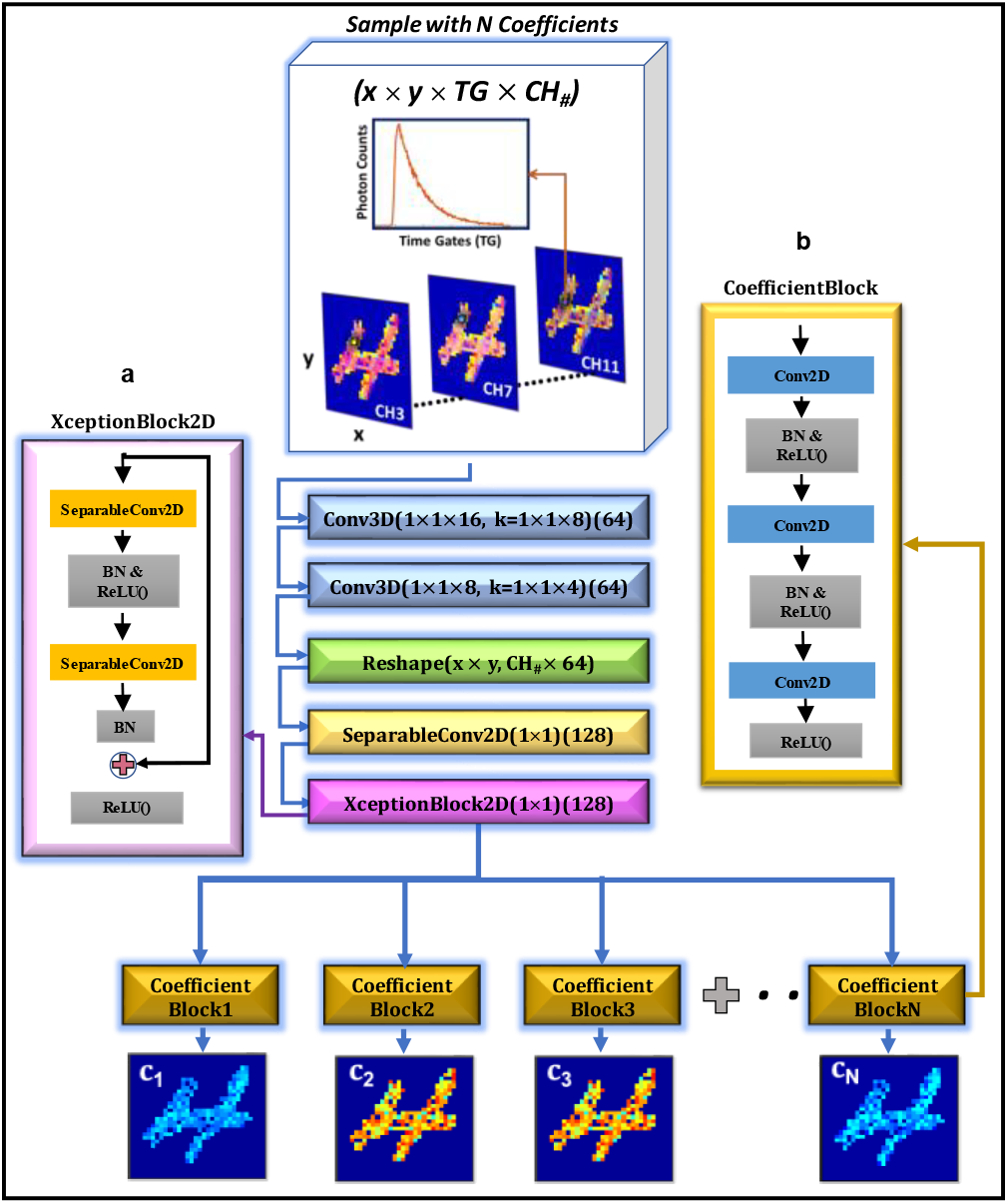
UNMIX-ME model architecture. HMFLI input mapped to spatially independent unmixed fluorescence coefficient values (**a**) “XceptionBlock” [7] comprised of 1×1 separable convolutions [8]. (**b**) “CoefficientBlock” structure.

As Fig. 2 illustrates, the data generation workflow for efficient but comprehensive training employed in this work partially follows the scheme of our previous work. [6], [11] In brief, a binary handwritten number dataset EMNIST was used for assignment of spatially-independent random variables of TPSF (*Γ*(*t*)) as provided in **Eq. (1)**. These variables include the lifetime values of fluorescent species involved (*τ*_*n*_), associated relative abundance coefficients (*c*^*n*^) and intensity scalar (*I*, expressed in photon counts). Note that these abundance coefficients are the output retrieval of UNMIX-ME (i.e. *c*_1_, *c*_2_ and *c*_3_ in the case of a tri-exponential application, **Eq. (2)**) [1]). Additionally, the instrument response function, *IRF*^*λ*^(*t*), is considered in the simulations to replicate as closely as possible experimental settings (example for three molecular species):

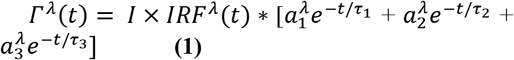

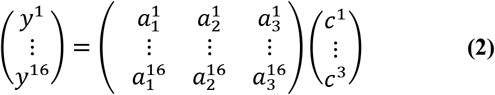

where *IRF*^*λ*^(*t*) corresponds to the instrument response function, 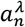 the relative spectral brightness of the nth fluorophore at the wavelength *λ*, and *I* to the overall photon counts to be detected. All variables used during spatially-independent generation of TPSFs were assigned at random value over wide bounds (ex., Fig. 2: (*τ*_1_, *τ*_2_, *τ*_3_, *I*) ∈ [0.9-1.1 ns, 0.3-0.4 ns, 0.55-0.65 ns, 50-500 p.c.]). These bounds represent typical values in NIR fluorescence imaging. It is trivial to extend these expressions to include *n*_*coeff*_ > 3 for both the CNN and the simulation workflow (example given in **Fig. S1** for *n* = 4).

**Fig 2.**
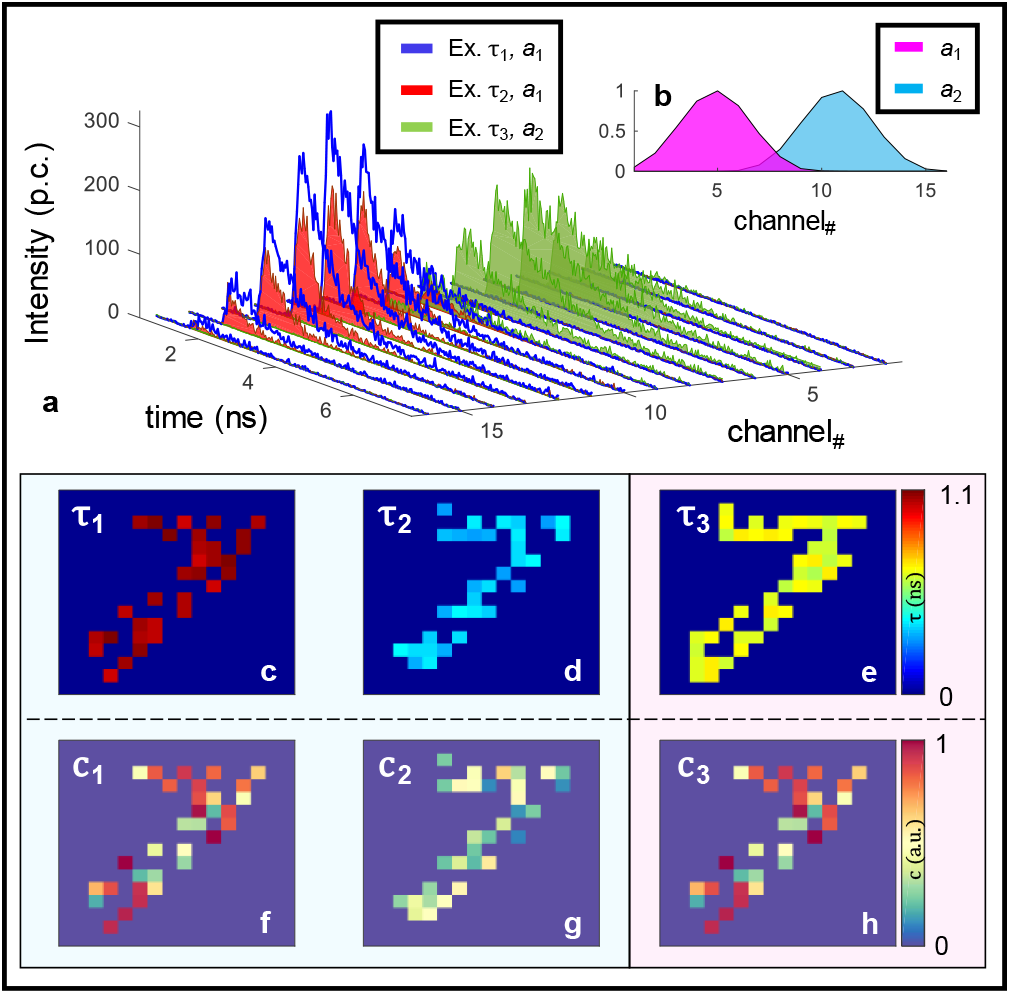
Data simulation workflow. A binary MNIST image is assigned lifetime values within three bounds (**c-e**). Using these values, along with spatially unique spectra (average given in **b**) for gathering intensity multipliers, 16 TPSFs are created at each non-zero spatial pixel (**a**). The coefficients are calculated shortly after (**f-h**).

Fig. 2a illustrates an example case of tri-specie unmixing in the challenging case of two fluorophores possessing the same emission spectral profile – a phenomenon inherent to both endogenous and FRET imaging. Accurate retrieval of abundance coefficients from fluorescent species possessing similar emission profiles necessitates either complex and heavily restrictive imaging protocols or time-consuming analytic pipelines based around iterative fitting. To ensure UNMIX-ME was sensitive to spectral bleedthrough, each spatial location which did not possess all three species was made to map all non-present coefficient values to zero. Greater detail regarding the simulation workflow is contained in the *Supplementary Materials Section 1* and GitHub repository [12]. Thus, each simulated 4D dataset was of size 16×16×256×16 (*x*, *y*, time-points, wavelength channels). The presented network model was built using *Tensorflow* with *Keras* backend in *python*. Training was performed over approximately 100 epochs using a total number of 2,500 datasets (80%-20% training-validation) and mean-squared error (MSE) loss. The total time for data simulation and training was 40 minutes and 15 minutes, respectively (NVIDIA Titan GPU). Further details of the training curves per number of species and t-distributed Stochastic Neighbor Embedding (t-SNE) visualizations of the different training coefficient ranges across the network are displayed in **Fig. S2**.

First, 250 tri-specie (two spectra, three lifetime) spectral TPSF data were simulated to illustrate how both our DNN and non-linear spectral unmixing coupled with least-squared bi-exponential lifetime fitting (LSQ+F) perform versus ground-truth during tri-spectral unmixing *in silico* (Fig. 3). Two overlapping gaussian profiles (Fig. 3a) were used to mimic independent 16-channel emission spectra. Further, lifetime parameters were assigned at random between three set bounds: (*τ*_1_, *τ*_2_, *τ*_3_) ∈ [0.95-1.1, 0.3-0.45, 0.55-0.7] ns. 2,500 separate data were generated for model training. Fig. 3(e-g) illustrates high spatial concordance with regards to all three coefficients, which is confirmed by the high SSIM values listed in Fig. 3k. Though high SSIM values are observed through LSQ+F retrieval of *c*_1_, a dip in accuracy is observed for both remaining coefficients. This dip is accuracy is not surprising given that *c*_2_ and *c*_3_ possess the same emission profile and depended upon often error-prone, iterative lifetime fitting to correct the coefficient value obtained through spectral decomposition. **Fig. S3** provides an example of two-spectra, four-specie unmixing which further supports this observation. For experimental validation, coefficient values were obtained from the HMFLI time domain reconstruction of a NIR-FRET well-plate with varying volumetric fractions of Transferrin (Tf) conjugated AF700 and AF750 dye, as illustrated in Fig. 4. The 4D input dataset was of size 64×64×256×16. The time domain reconstruction process is further explained in *Supplementary Section 2*. Förster Resonance Energy Transfer (FRET) unmixing is a uniquely complex two-spectra three-specie problem given the under and overestimation of the donor and acceptor fluorescent contribution due to quenching, respectively. [13], [14] Though, this effect was easily taken into account during data simulation (detailed in *Supplementary Section 1*). [15] The UNMIX-ME framework allowed for retrieval of total Tf-AF700 and Tf-AF750 coefficient values adhering much more closely to the expected values Fig. 4(p-t) compared to conventional LSQ+F, Fig. 4(f-j). All wells containing Tf-AF700 (*c*_1*T*_) were prepared with constant volume and thus the decreasing trend observed through LSQ+F (Fig. 4h) is much higher than that illustrated through UNMIX-ME (Fig. 4r). Further, the second and third row were both prepared with same increasing volumes of AF750 (*c*_2_), and thus the *c*_2_ trend observed should be identical. Fig. 4i illustrates an overestimation of Tf-AF750 in the second row via LSQ+F – an expected result given the overestimation of acceptor concentration in the case of FRET. In contrast to this, UNMIX-ME provides *c*_2_ quantification with significantly higher overlap (Fig. 4s). Moreover, though FRET quantification (FD (%)) through both UNMIX-ME (Fig. 4t) and LSQ+F (Fig. 4j) were relatively similar for the 1:1 to 3:1 cases, the single-specie well (0:1) was overestimated through iterative fitting while UNMIX-ME correctly estimated values of zero across the entire well. The quenched donor (*c*_1_^∗^) abundance estimated through UNMIX-ME (Fig. 4q) provides both a much more easily distinguishable increasing trend from well-to-well (as expected) and correctly assigned values of zero at the 0:1 ROI than when using LSQ+F (Fig. 4g).

**Fig 3.**
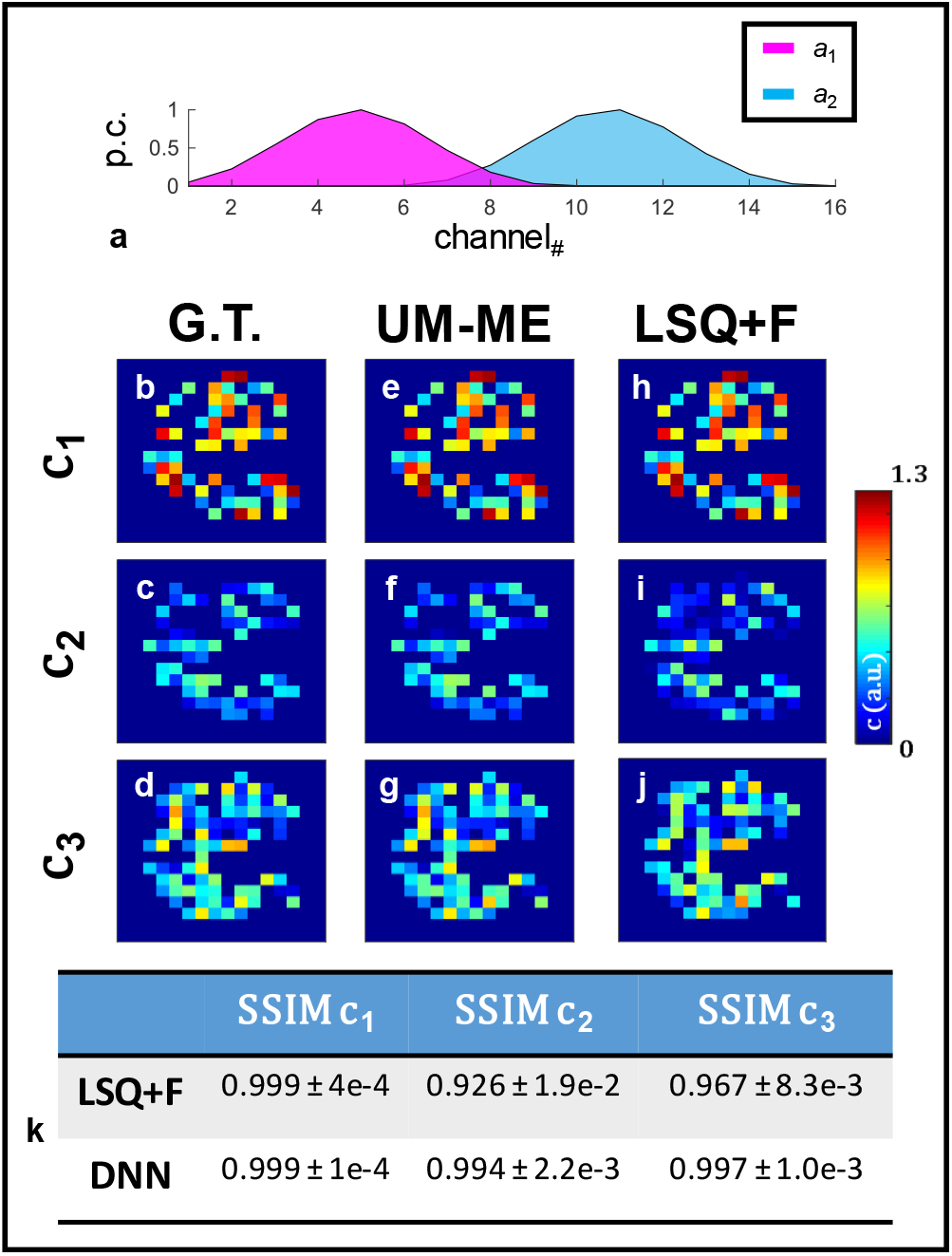
Three-coefficient spectral unmixing *in silico*. Averaged spectra used for simulation (**a**) are given. (**b-d**) Ground-truth values are illustrated as well as the coefficients retrieved via DNN lifetime unmixing (**e-g**) and conventional LSQ+F fitting (**h-j**). **Table S1** provides average and standard deviation MSE values calculated across 100 test samples as for additional performance quantification.

**Figure 4.**
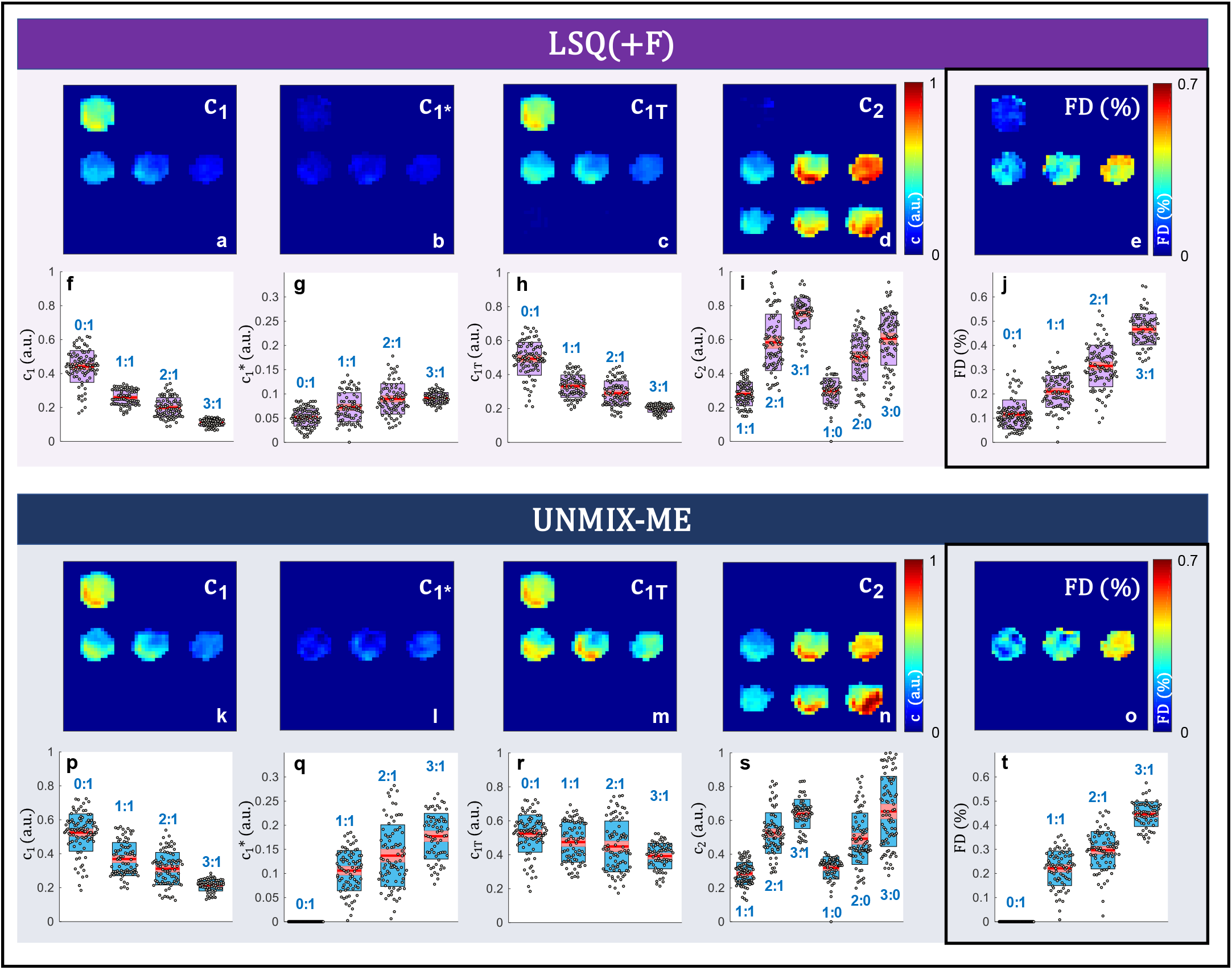
HMFLI-FRET *in vitro*. Results from non-linear iterative spectral decomposition combined with lifetime fitting (LSQ+F) (**a-j**) and UNMIX-ME (**d-t**) are given. Boxplots of coefficient values retrieved at each ROI (labeled by acceptor/donor ratio) are given per reconstruction.

Finally, UNMIX-ME was used for two complex cases of HMFLI-FRET imaging *in vivo*. MFLI allows to quantitatively report on target-receptor interaction via *in vivo* lifetime-based FRET and further FD (%) quantification [16], [17]. First, HMFLI data acquired from a nude athymic mouse 6-hours post-injection with Tf-conjugated AF700 and AF750 FRET pair (in a 2:1, acceptor-to-donor ratio) was unmixed via both methods for comparison. The engagement of Transferrin (Tf) receptors in the liver allows for a change in lifetime and FD (%) compared to excretion organs like the bladder, therefore allowing for further organ classification. [13] For this task and as previously shown for *in vitro* samples the FD (%) will be resolved for both methods from the resulting unmixed abundance coefficients of the unquenched and quenched donor (*c*_1_ and *c*_1_^∗^ respectively). Ideally, *c*_2_/*c*_1 *total*_) ratios closely correspond with the 2:1 injected acceptor to donor concentration. Furthermore, the resolved coefficients should reflect the difference in lifetime and FRET between liver and bladder organs. For brevity, the results of this experiment are illustrated in **Fig. S4** and further discussed in *Supplementary Section 4*. UNMIX-ME exhibited the capability to better resolve the change in lifetime and FD (%) upon the retrieved donor (*c*_1 *total*_) and acceptor (*c*_2_) coefficients compared to LSQ(+F). UNMIX-ME better depicted the expected quenching of the donor in the liver which contains a high volume of Tf receptors but not in the bladder. At the same time, the injected 2 (***c***_**2**_) to 1 (***c***_**1 *total***_) acceptor to donor concentration was better described by UNMIX-ME’s unmixed relative abundance coefficients. The second and final *in vivo* case, results of which are displayed in Fig. 5, involves 76-hours post-injection data acquired from a nude athymic mouse bearing a tumor xenograft and injected with HER2 targeted-Trastuzumab AF700 and AF750 FRET pair (in a 2:1, acceptor-to-donor ratio). TZM is used to treat metastatic breast cancer in the clinic and has been recently proposed for fluorescence lifetime imaging. [18] Engagement to HER2 receptors present in the tumor region will result in a decrease in lifetime and increase in FRET interaction (FD (%)). These expected changes should be also reflected in the unmixed acceptor, quenched/non-quenched donor coefficients retrieved. As for the previous FRET in vivo experiment, the FD (%) will be retrieved via both LSQ(+F) and UNMIX-ME. HMFLI data was reconstructed for a 128×128 resolution and acquired over 6 minutes for 16 detection wavelengths (channels) through high data compression as highlighted elsewhere [5]. Thus, the 4D input dataset was of size 128×128×256×16. The tri-component (unquenched donor + quenched donor (*c*_1 *total*_) and acceptor (*c*_2_) spectral lifetime unmixing was retrieved using both LSQ+F Fig. 5(d-f) and UNMIX-ME in Fig. 5(a-c). Quantifications for these regions are displayed in Fig. 5(g-i). *In vivo* spectral unmixing results illustrated in Fig. 5 were obtained through photon-count thresholding to focus solely on the xenograft region. The FRET-FD (%) quantification obtained through both methods is in high concordance – a result previously observed at higher FRET levels *in vitro* (Fig. 4(j, t)). However, the comparison of results of acceptor *c*_2_ in Fig. 5h to donor *c*_1 *total*_ in Fig. 5g illustrate that *c*_2_/*c*_1 *total*_ratios obtained through UNMIX-ME correspond much more closely with the 2:1 injected acceptor to donor concentration than those obtained through LSQ+F (further supported in **Fig. S6**).

**Fig 5.**
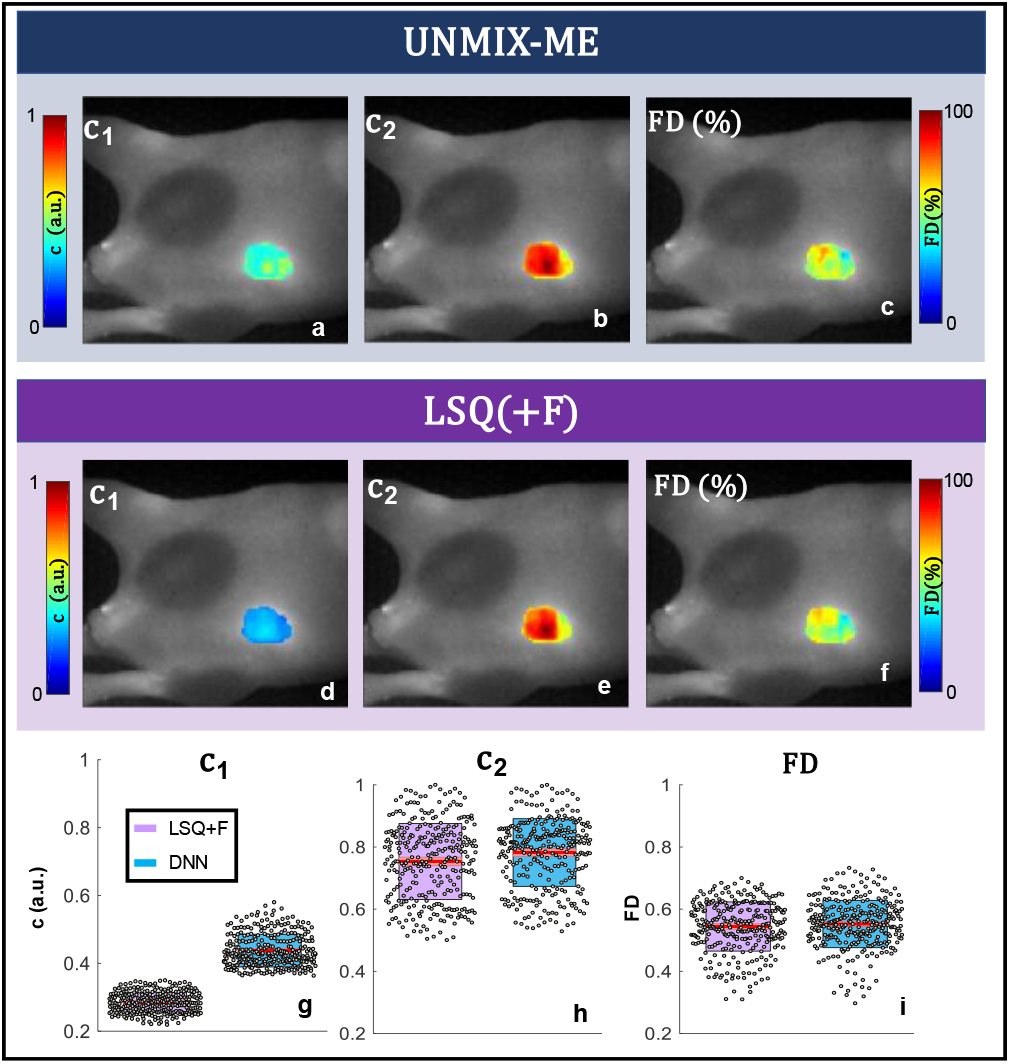
HMFLI-FRET imaging of tumor xenograft *in vivo*. Mouse xenograft imaged at 76-hours post-injection of Trastuzumab (AF700 & AF750). Results from UNMIX-ME (**a-c**) and LSQ+F (**d-f**). Boxplots are given for further quantitative clarity (**g-i**).

In summary, we present UNMIX-ME, a DNN-based workflow trained with simulation data for fluorescence lifetime unmixing. Furthermore, unlike intensity-based approaches paired with iterative lifetime fitting, UNMIX-ME simultaneously leverages intensity and lifetime features for sensitive retrieval of relative abundance coefficients of the mixed fluorophores. UNMIX-ME provided improvement over a conventional sequential iterative fitting methodology by demonstrating higher concordance with ground truth during tri- and quadri-abundance coefficient retrieval (**Fig. S1**). Furthermore, UNMIX-ME provided accurate unmixed profiles for complex HMFLI data from FRET interactions *in vitro* and for two independent non-invasive HMFLI *in vivo* experiments, where the retrieved abundance donor and acceptor coefficients were used to calculate FD (%) and further validate the injected 2:1 acceptor to donor concentration ratio of the dyes. Through the calculated FD (%) UNMIX-ME effectively accomplished organ classification for Transferrin based targeting of liver and bladder and tumor xenograft targeting of HER2 receptors through Trastuzumab. Thus, UNMIX-ME unlocks the possibility for efficient and accurate fluorescence lifetime unmixing of multiplexed data at the macroscopic level. Moreover, it illustrates the future potential of applying DNNs for microscopic unmixing.

## Acknowledgments

We gratefully acknowledge the support of NVIDIA Corporation with the donation of the Titan Xp GPU. We would like also to acknowledge the support of Drs. Margarida Barroso and Alena Rudkouskaya in providing the animal model as well as the conjugated probes. This work was supported by the National Institutes of Health Grants R01 EB19443, R01 CA207725 and R01 CA237267.

## Notes

#### Summary of Updates

The following major updates have been made: - Additional experiments added. - Proper FRET unmixing (quenched and unquenched donor specie separation). - Deeper probing into DNN-based metrics and interpretation.

