## Supplementary Materials for "Spectral and lifetime fluorescence unmixing via deep learning"

Supplementary Information
for

### Spectral and lifetime fluorescence unmixing via deep learning

Jason T. Smith[+], Marien Ochoa[+] and Xavier R. M. Intes[*]

### 1. Data Simulation Workflow

##### Simulating Application-Specific Hyperspectral FLI Data

As was discussed briefly, the simulation routine developed for this work, which has been validated only for HMFLI unmixing to date, is detailed in great depth on our *GitHub* repository https://github.com/jasontsmith2718/UNMIX-ME (along with corresponding data, relevant script, etc.), but will be described in brief here.

In order to generate experimentally realistic hyperspectral FLI data through this approach, one must have the following:

- Pure emission spectral profiles of each fluorescent specie comprising the sample.

- Each fluorophore's lifetime (known within +/- 100 ps).

- Instrument Response Function $IRF^{\lambda}(t)$ of the HFLI apparatus.

At the start of every simulation iteration, an MNIST (Modified National Institute of Standards and Technology database) binary figure is chosen at random. This figure is subsequently downsampled by a factor of two (32×32 -> 16×16) and used for subsequent value assignment. At every non-zero pixel, values of lifetime (τ) are assigned at random – with the $n_{\tau} = n_{coeff}$ (example for four-lifetime case illustrated in **Figure S1**). Each pixel is also assigned individual spectral profiles (all initially max-normalized) which are immediately multiplied by a random scalar value between 50-500 and assigned to specie-specific pre-allocated arrays of size $(x \times y \times n_{channels})$. Next, performing a loop over all non-zero pixels, a random value between zero and one is obtained in order to select a combination of fluorescent species to include (the total number of which is always equal to $n_{coeff}$!). Afterwards, a nested loop generates TPSFs ($\Gamma^{\lambda}(t)$) across all 16 channels via **Eq. S1**

$$\Gamma^{\lambda}(t) = I \times IRF^{\lambda}(t) * e^{-t/\tau_n} \qquad \textbf{(S1)}$$

Where $I$ indicated the scalar intensity value of the *n*th fluorophore at each channel. This is performed *n* times to account for all fluorophores, simulating a total of n TPSFs – each of which undergo a collective sum (including all *n* TPSFs) followed by Poisson noise assignment. To account for potential system-dependent laser jitter, randomly selected TPSFs are shifted around their final position (+/- three integer values). This 16-TPSF generation repeats for all non-zero values across the image. Concurrently, values of one are assigned to an array initialized with all zeros of size $(x \times y \times n_{coeff})$ to indicate the fluorophores present at each pixel.

Next, ground-truth (G.T.) coefficient values are obtained (example illustration shown **Fig. S1(h-k)**), possessing size $(x \times y \times n_{coeff})$. By summing over the fourth dimension of the 16-channel mono/multi-specie TPSF data, continuous wave (CW) data possessing size $(x \times y \times n_{channels})$ were calculated. Afterwards, iterative intensity-based spectral unmixing was performed at each non-zero-pixel location via **Eq. 1** using with each pure emission spectral profile (max-normalized ex. illustrated in **Fig S1a**) to obtain values for $c$. For the case of multiple species possessing the same emission profile, the coefficient value obtained through the simplistic intensity-based technique is further multiplied by a ratio of the maximum value of each specie's initially assigned pre-allocated spectral profile over the sum of all maximum intensity values possessing the same emission profile (a maximum of two values per spectra in this report). Finally, to obtain true G.T. values and ensure that the network is trained to map spectral bleed-through to zero, the array of obtained coefficient values is multiplied by the binary array used to keep track of fluorophore presence at each $(x, y)$ location. This single matrix dot product zeros out all non-contributing coefficient values and is the reason behind the sparsity illustrated in **Fig. 2**, **Fig. S2** and **Fig. S2**.

##### Correction in the Case of Resonance Energy Transfer

The nonradiative RET phenomena occurs when two conditions are satisfied: **1)** the emission and excitation spectra of two fluorescent species overlap, **2)** both fluorophores are oriented in a particular fashion relative to each other and are in close proximity (within < 10nm). Spectral unmixing of fluorescent species undergoing Resonance Energy Transfer (RET) is a particularly special case which necessitates correction via fluorophore-specific photophysical values to obtain accurate quantification. Correcting for RET during unmixing has been conventionally performed using the following expression (**Eq. S2**)[1].

$$a_T = c_D \times (1 - E) \times a_D + a_A \times c_A + c_D \times E \times (\phi_A/\phi_D) \times k(\lambda) \times a_A \qquad \textbf{(S2)}$$

Where $a$, c, $\phi$, E and $k(\lambda)$ correspond to the spectral profile, abundance coefficient, quantum yield (with T, D and A notating total, donor and acceptor), FRET efficiency and the ratio of pure donor over pure acceptor emission spectra, respectively. The spectra obtained for calculating $k(\lambda)$ were those collected from the wells containing only AF700 and AF750 during the well-plate HMFLI-FRET acquisition (**Fig. 4**).

By **Eq. S2**, conventional linear unmixing methods will underestimate the value of $c_D$ (by a factor of (1-E) and overestimate $c_A$ by a factor of $c_D \times E \times (\phi_A/\phi_D) \times k(\lambda)$ – the results of which were observed in both **Fig. 4** and **Fig. 5** using LSQ+F. Thus, during the step of obtaining the fractional abundance coefficient, these values (either quickly found in literature [$E, \phi_A$, $\phi_D$])[2] or calculated directly ($k(\lambda)$)) for the AF700/AF750 NIR-FRET pair were used for abundance coefficient correction. By training UNMIX-ME to map directly to RET-corrected $AF700_{FRET}$ ($\tau \cong$ 0.3-0.45)ns, $AF700_{non\text{-}FRET}$ ($\tau \cong$ 0.95-1.0)ns and AF750 ($\tau \cong$ 0.5-0.65)ns, the spectral lifetime unmixing results presented herein were obtained. Given that the model presented herein utilized NIR fluorescent species possessing analytically demanding sub-nanosecond lifetimes, adaptation of UNMIX-ME for commonly used visible FRET pairs should be trivial.

#### 2. Time Domain Reconstruction from Single Pixel CS-HMFLI:

The reconstruction process from single pixel data acquisitions to time domain (TD) reconstructions is accomplished through the inverse solving of: ***M(t) = P x(t)*** for the image ***x(t)*** of the sample plane across 256 time points, where ***M(t)*** represents the raw fluorescence decay profiles acquired per pattern ***P***. The pattern's matrix serves to calculate the corresponding weights each measured pattern has for the sample plane image. Since the pattern's matrix ***P*** and time domain measurement matrix ***M(t)*** are known, then through the use of TVAL3 [3] inverse solver, the fluorescence decay profiles per each pixel of the image sample plane, can be retrieved. To obtain continuous wave intensity images, the representations across the 256 time points can be added. In order to obtain lifetime reconstructions, each of the fluorescence decay's per pixel are bi-exponential fitted through a least-squares based minimization algorithm. The bi-exponential mean lifetime model is represented by: $\boldsymbol{(A_1/100) \times \tau_1 + (A_2/100) \times \tau_2}$ where A's represent the FRETing Donor and Acceptor fraction and tau's their respective lifetime values.

#### 3. UNMIX-ME DNN Model: Relevant Metrics and Interpretation

**Fig. S2a** illustrates the validation MSE loss over 100 epochs averaged across five separate training cycles. Notably, the MSE loss of $c_1$ (green) converged to a much lower value other two (red and blue). This observation comes without surprise given that the retrieval of $c_1$ was obtained primarily through intensity-based unmixing in immense contrast to both $c_2$ and $c_3$ which share the same emission profile and both possess sub-nanosecond lifetimes differing by an average of just 0.3 ns.

Further, **Figure S2b** illustrates the validation MSE loss over 90 epochs averaged across five separate training cycles for a two-spectra, four-specie case (results shown in **Fig. S2**). Comparatively, not a single loss value gets anywhere near the previously mentioned lowest MSE obtained through **Fig. S2a** Since, in this case there would be no way to perform spectral unmixing via intensity alone, this observation supports one's intuition that UNMIX-ME would experience greater difficulty in learning the inverse mapping in a much more complicated scenario. Further, the MSE validation loss for both $c_3$ and $c_4$ is significantly lower than the MSE of $c_1$ and $c_2$. Though both spectral profiles are similar (evidenced by **Fig S1a**), the lifetime values for the ($c_1$ , $c_2$) pair, ([0.5-0.6] ns, ([1.0-1.1] ns) is significantly closer than that of the ($c_3$ , $c_4$) pair ([0.25-0.35] ns, ([1.5-1.6] ns) and thus the inverse mapping is understandably more challenging to undertake for the first pair than the second. This is further supported by **Fig S3(m, n)** which present stark differences in both SSIM and MSE performances between the fluorescent pairs. Though, in all cases presented herein, UNMIX-ME performs exceedingly well.

**Fig S2(c-f)** provides the t-distributed Stochastic Neighbor Embedding (tSNE)-reduced 3D projection of flattened DNN activations extracted at the mid-way point within each of the four separate branches during the forward-pass of 1,000 test datasets. These data, generated with the same two-spectra/four-specie parameters as those employed in **Fig S1/S3**, were each assigned just a single spectral pair and lifetime quartet at random instead of spatially-independent assignment. As can be observed from each coefficient's tSNE scatter plot (where each sphere represents a simulation data voxel) the color gradients, which indicate ground-truth mean-abundance values for both pairs, show clear continuity in all four cases. This provides further insight into the network's capability for extracting these complex spectral and temporal features during inference.

#### 4. *In vivo* HMFLI-FRET: Supplementary Experiment – Transferrin/Transferrin Receptor (Tf/TfR) Engagement

HMFLI data was acquired from a nude athymic mouse after 6-hours post-injection with conjugated Transferrin (Tf) AF700 and AF750 FRET pair (in a 2:1, acceptor-to-donor ratio). Ideally the 2:1 acceptor to donor ratio must be represented by the abundance coefficients retrieved by UNMIX-ME or LSQ(+F). Previous *in vivo* work in mice has demonstrated the ability to characterize FRET engagement through conjugated Tf-probes in the liver and compare against the lifetime on the bladder [4]. The liver is known to display a high density of Transferrin receptors and thus possesses correspondingly high degrees of FRET when imaged [5]. However, the urinary bladder acts solely as an excretion organ, eventually accumulating free dye – possessing negligible potential for FRET occurrence. HMFLI data was acquired for over 12 minutes for a 64x64 resolution for 16 detection wavelengths (channels), using only 50% of the total required single pixel measurements. Therefore, resulting in a 4D input dataset of size 64x64x256x16.

**Fig. S4** reports the abundance coefficients and FRET percentage retrieved through both UNMIX-ME and LSQ+F (**Fig. S4(a-d)** and **Fig. S4(e-h)**, respectively.) As previously encountered *in vitro* (**Fig. 4**) the LSQ+F methodology underestimates the AF700 abundance coefficient ($c_1$) in the liver (**Fig. S4f**, FRET site), overestimates the AF750 abundance ($c_2$) in the bladder (**Fig S4g**, non-FRET site) and provides a non-zero FRET percentage across the urinary bladder in contrast to conventional knowledge. The FRET percentage retrieved at the liver ROI is in high agreement across both methods, as was also demonstrated previously at high-FRET wells (**Fig. 4(j, t)**). However, UNMIX-ME provides a total AF700 abundance in higher concordance with levels expected (**Fig. S4(a, b)**), FRET levels of zero in the urinary bladder (**Fig. S4d**) and an AF750 abundance trend that matches the expected 2:1 ratio closely across the liver ROI (**Fig. S4g**).

| | MSE $c_1$ | MSE $c_2$ | MSE $c_3$ |
|---|---|---|---|
| LSQ+F | 4.2e-5 ± 9.0e-6 | 3.7e-3 ± 1.1e-3 | 3.9e-3 ± 1.1e-3 |
| DNN | 2.3e-5 ± 7.6e-6 | 3.1e-4 ± 1.1e-4 | 3.2e-4 ± 1.0e-4 |

**Table S1**. Mean-squared error calculated for all three coefficient values through both techniques (complement to **Fig. 3.**).

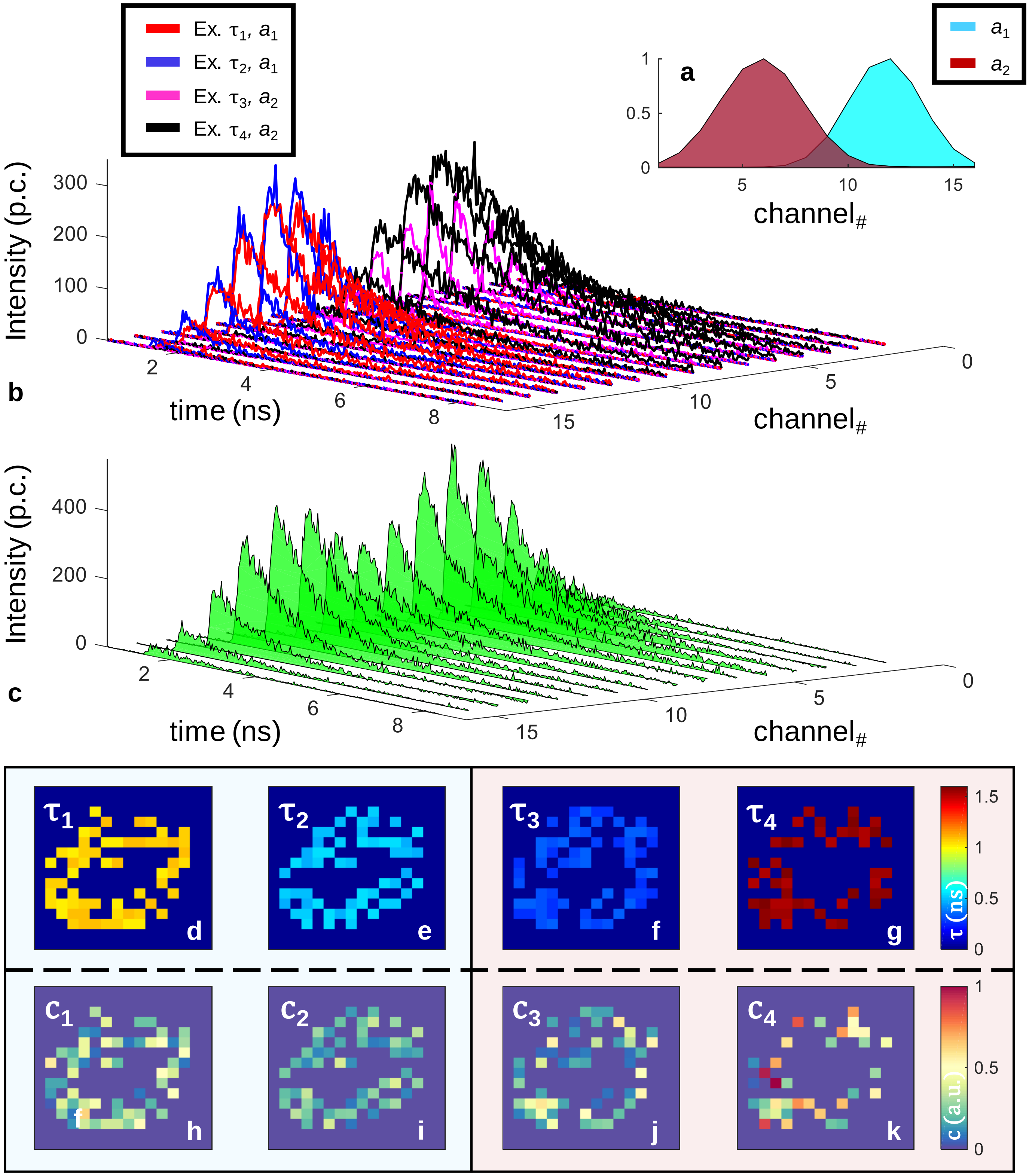


**Figure S1**. Information relevant for a two-spectra (**a**), four-specie (**d-g**) in silico spectral lifetime unmixing performance assessment. Example mono-specie HFLI data (b) along with an example containing a mix of all four (c) is given for illustrative purpose. Blue and red boxes indicate the spectra (**a**) to which each of the lifetime and coefficient (**h-k**) values belong.

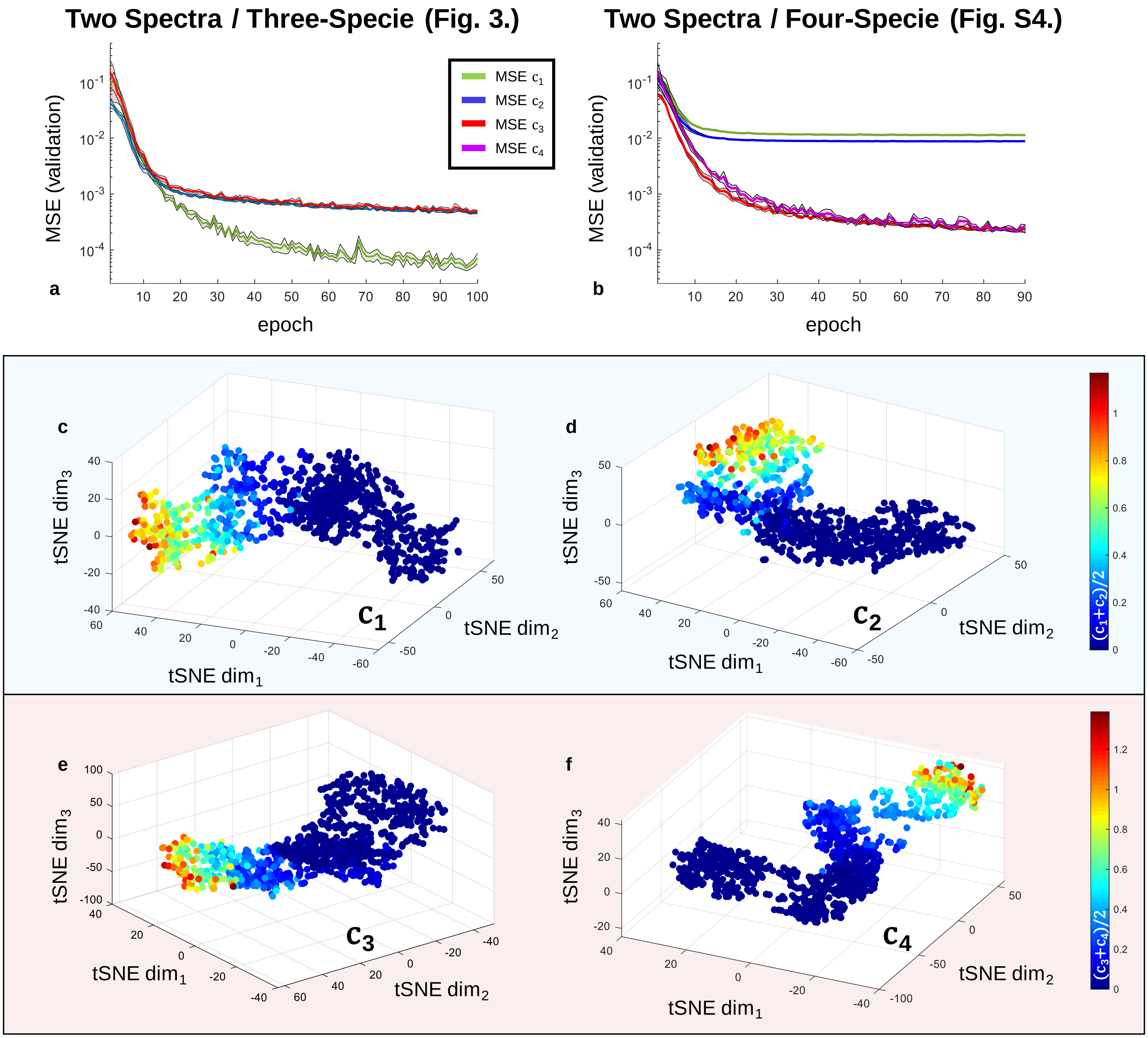


**Figure S2**. Metrics relevant to the UNMIX-ME deep convolutional neural network. The MSE validation loss curves were obtained for each coefficient separately over five separate training iterations for two separate cases - two-spectra, three-lifetime (**Fig. 3**) and two-spectra, four-lifetime (**Fig. S4**). The average and standard deviation of each is provided (**a**, **b**). Four the two-spectra/four-specie case, 1,000 separate samples were generated and fed into the network in order to perform a tSNE assessment of each branch individually (**c-f**).

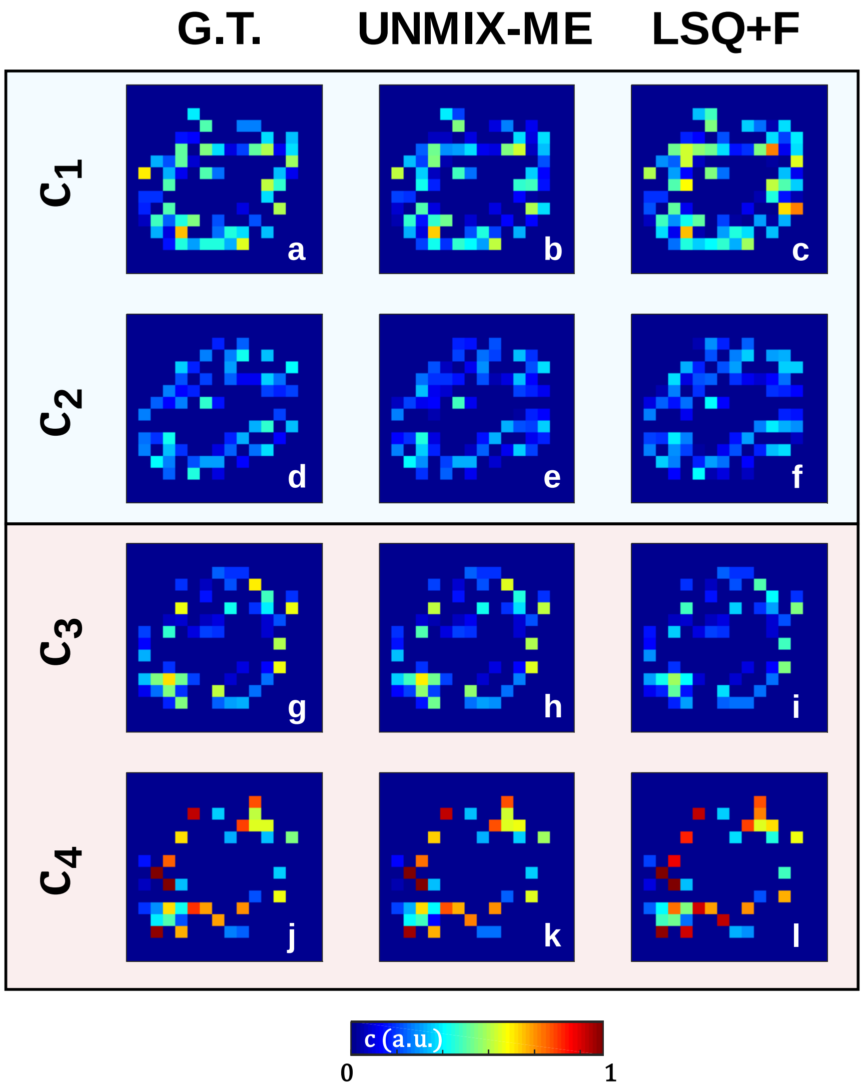


| | SSIM $c_1$ | SSIM $c_2$ | SSIM $c_3$ | SSIM $c_4$ |
|---|---|---|---|---|
| **LSQ+F** | 0.826 ± 8.1e-2 | 0.837 ± 7.4e-2 | 0.955 ± 6.4e-2 | 0.977 ± 6.4e-2 |
| **DNN** | 0.924 ± 3.0e-2 | 0.897 ± 3.5e-2 | 0.997 ± 1.5e-3 | 0.999 ± 6.0e-4 |

**m**

| | MSE $c_1$ | MSE $c_2$ | MSE $c_3$ | MSE $c_4$ |
|---|---|---|---|---|
| **LSQ+F** | 3.58e-2 ± 1.6e-2 | 1.52e-2 ± 6.2e-3 | 4.01e-3 ± 4.7e-3 | 4.84e-3 ± 1.6e-2 |
| **DNN** | 1.17e-2 ± 4.6e-3 | 8.81e-3 ± 2.9e-3 | 2.42e-4 ± 1.0e-3 | 1.99e-4 ± 1.0e-5 |

**n**

**Figure S3**. Spectral lifetime unmixing performance obtained via both LSQ+F and UNMIX-ME versus ground-truth. A single illustration is given for qualitative assessment. Both SSIM (**m**) and MSE (**n**) were calculated for 250 test samples to provide quantification with regards to image-to-image similarity and direct one-to-one value comparison.

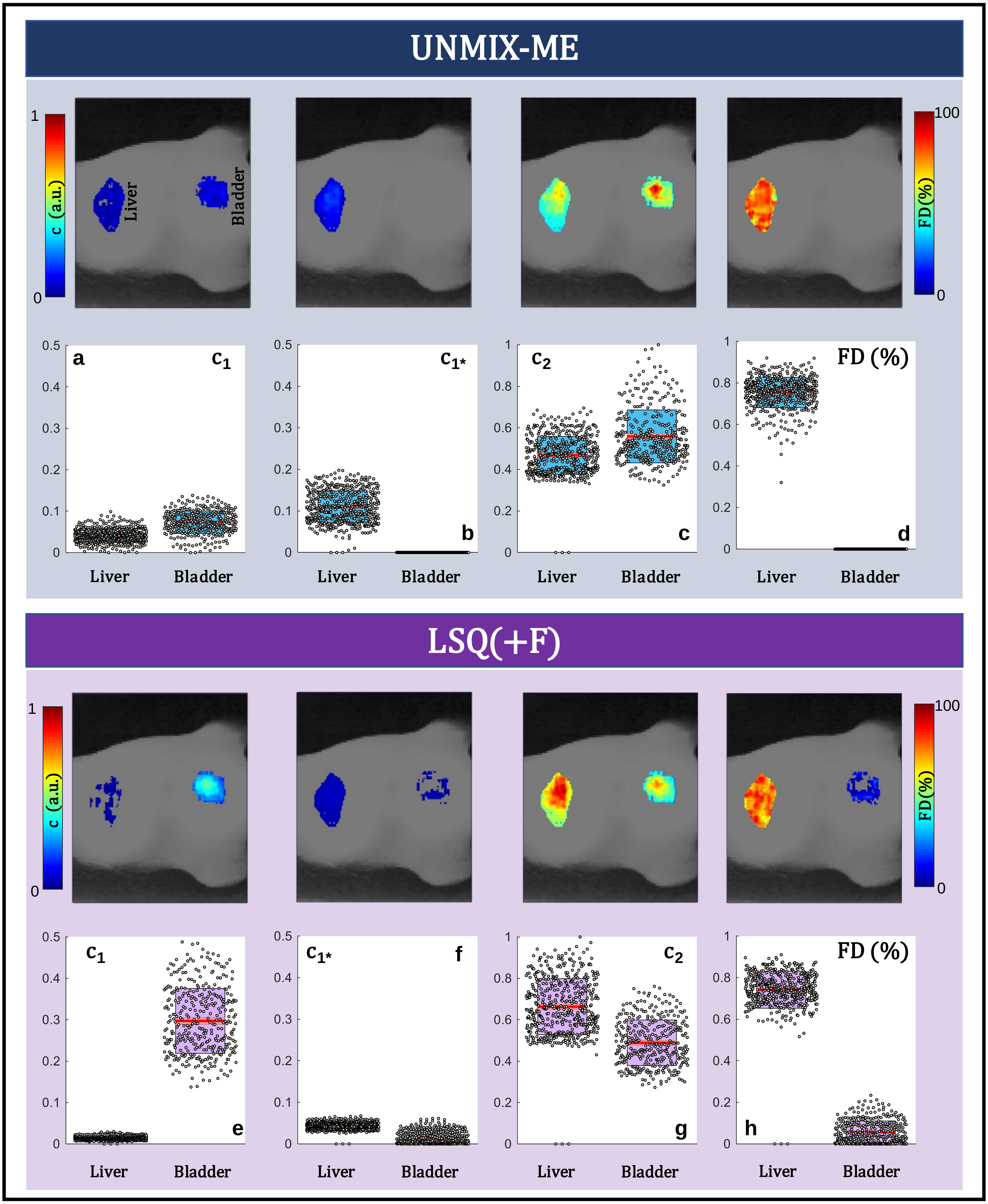


**Figure S4**. HMFLI-FRET imaging of Transferrin receptor (TfR) engagement *in vivo*. Results from UNMIX-ME (**a-d**) and LSQ+F (**e-h**) are provided. Resolved liver and bladder areas are displayed per reconstruction.

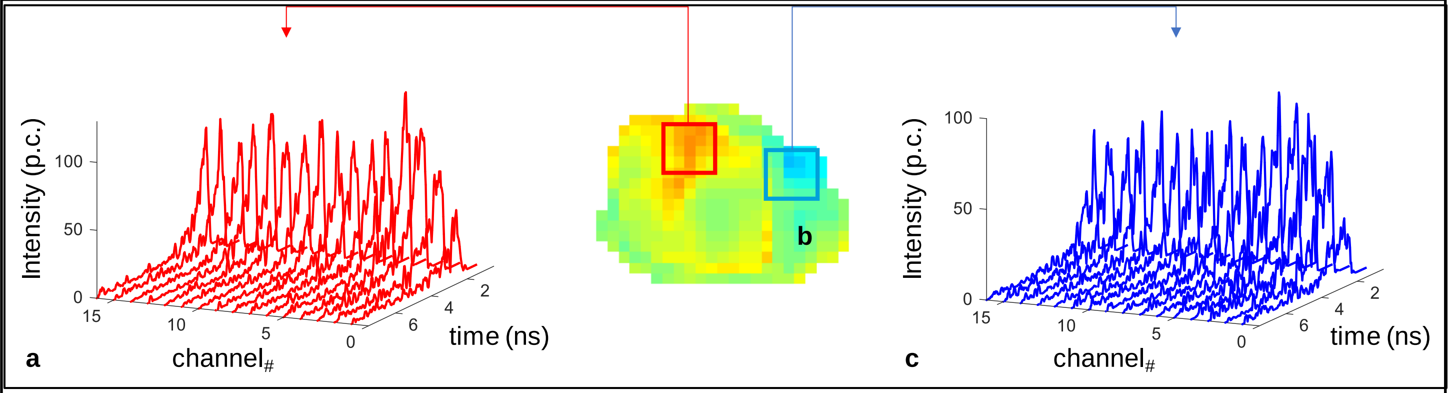


**Figure. S5**. Averaged HMFLI TPSF data at two different regions of the tumor xenograft (**b**). The regions were chosen due to their high (**a**) and low (**c**) FRET quantification.

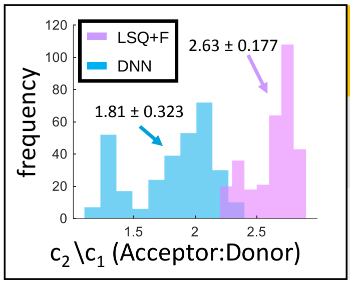


| | $c_1$ (a.u.) | | $c_2$ (a.u.) | | FRET (%) | |
|---|---|---|---|---|---|---|
| **LSQ+F** | 0.285 ± 3.0e-2 | 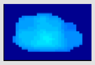 | 0.753 ± 0.12 | 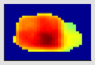 | 54.4 ± 7.91 | 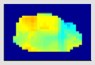 |
| **DNN** | 0.438 ± 4.8e-2 | 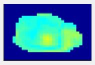 | 0.784 ± 0.11 | 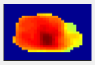 | 55.2 ± 7.62 | 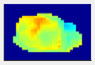 |

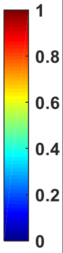


**Figure S6**. Additional information provided to complement the in vivo spectral lifetime quantification performed in **Fig. 5**. Histogram provides all acceptor/donor ratios obtained for each spatial location across the tumor ROI. The table provides averaged and standard deviation values for both coefficients and the FRET % values obtained through both methods.
